# Wisdom of artificial crowds feature selection in untargeted metabolomics: An application to the development of a blood-based diagnostic test for thrombotic myocardial infarction

**DOI:** 10.1101/165977

**Authors:** Patrick J. Trainor, Roman V. Yampolskiy, Andrew P. DeFilippis

## Abstract

**Introduction:** Heart disease remains a leading cause of global mortality. While acute myocardial infarction (colloquially: heart attack), has multiple proximate causes, proximate etiology cannot be determined by a blood-based diagnostic test. We enrolled a suitable patient cohort and conducted an untargeted quantification of plasma metabolites by mass spectrometry for developing a test that can differentiate between thrombotic MI, non-thrombotic MI, and stable disease. A significant challenge in developing such a diagnostic test is solving the NP-hard problem of feature selection for constructing an optimal statistical classifier.

**Objective:** We employed a Wisdom of Artificial Crowds (WoAC) strategy for solving the feature selection problem and evaluated the accuracy and parsimony of downstream classifiers in comparison with embedded feature selection via the Lasso and Elastic Net.

**Materials and Methods:** Artificial Crowd Wisdom was generated via aggregation of the best solutions from independent and diverse genetic algorithm populations that were initialized with bootstrapping and a random subspaces constraint.

**Results / Conclusions:** WoAC feature selection performed favorably compared to Lasso and Elastic Net solutions. The classifier constructed following WoAC feature selection had a cross-validation estimated misclassification rate of 2.6% as compared to 26.3% via the Lasso and 18.5% via an Elastic Net. The classifier warrants further evaluation as a diagnostic test in an independent cohort.

## 1. Introduction

Heart disease remains the most prevalent cause of death worldwide despite dramatic reductions in the incidence of heart disease associated mortality in developed countries (Mozaffarian et al., 2016). Acute Myocardial Infarction (AMI), an acute manifestation of heart disease, is characterized by myocardial ischemia (oxygen starvation in heart muscle) and necrosis (a form of cell death) secondary to atherosclerotic plaque disruption or other cause. While ischemia and necrosis are the common pathological characteristics of AMI there are multiple underlying causes that can lead to ischemia and necrosis (Thygesen et al., 2012). An important etiological distinction can be made between thrombotic and non-thrombotic MI. Thrombotic MI results from spontaneous atherosclerotic plaque disruption (e.g. rupture or erosion) that results in the formation of an occluding coronary thrombus (Thygesen et al., 2012). In contrast, MI secondary to oxygen supply / demand mismatch resulting from conditions not associated with plaque rupture such as vasospasm or stress cardiomyopathy are categorized as non-thrombotic MI. These etiological distinctions are of critical importance as the course of treatment depends on the underlying cause and diagnostic misclassification may result in negative outcomes such as iatrogenic bleeding (Amsterdam et al., 2014).

Blood-based tests that measure the release of troponin from the myocardium can provide evidence of myocardial necrosis which may be used to substantiate a diagnosis of AMI (Newby et al., 2012). However, a non-invasive test that enables the discrimination of thrombotic from non-thrombotic MI has yet to be developed. In response, we set out to develop a blood-based test that could differentiate thrombotic MI from non-thrombotic MI and stable coronary artery disease (CAD). In developing a blood-based diagnostic test we chose a plasma medium—plasma contains enzymes, lipoproteins, hormones, metabolic intermediates, and other small molecules dissolved in suspension. Plasma provides a suitable medium as plasma is a repository of biochemicals that reflects the state of the entire organism at the time of sampling (Psychogios et al., 2011). We focused on metabolites—or more precisely small molecules—as metabolite concentrations are dynamic and reflect the “end result” of genetic factors, environmental exposures, and gene-environment interactions at the time of sampling (Nicholson et al., 2012). A patient cohort was recruited that allowed for the discrimination of thrombotic MI from multiple control populations (DeFilippis et al., 2015). This cohort consisted of three study groups: thrombotic MI, non-thrombotic MI, and stable coronary artery disease subjects. The non-thrombotic MI group controlled for metabolic changes associated with myocardial ischemia and necrosis, while the stable coronary artery disease group was used to control for the underlying disease process of atherosclerosis. Both control groups served as procedural controls as all three study groups underwent a cardiac catheterization procedure following study enrollment.

Whole blood was collected immediately prior to cardiac catheterization from subjects in each of the study groups and plasma metabolite relative abundances were determined by an un-targeted mass spectrometry approach. Two challenges arise in developing a diagnostic classifier from such data. The first is that the dimension of the feature space (1,032 detected metabolites) is significantly larger than the number of samples (11 thrombotic MI, 12 non-thrombotic MI, and 15 stable CAD). To determine the best subset of metabolites to be included in a classifier is thus a search problem for which a brute-force is not advisable. For example, to determine the optimal classifier with five metabolites, 9.7×10^12^ combinations are possible. Second, the parameter estimates of a classification model may be highly unstable given the small sample size. Consequently, an algorithm that searches for the best possible subset of metabolites for inclusion in a classifier should converge in reasonable time and should minimize a measure of expected prediction error. In addition to minimization of expected prediction error, a variable selection technique should seek to minimize the number of variables selected in this context given the prohibited cost of developing and validating targeted assays. In this paper, we evaluated the use of a Wisdom of Artificial Crowds (WoAC) (Yampolskiy & Barkouky, 2011) approach to variable selection for developing a blood-based diagnostic test for thrombotic myocardial infarction. A WoAC approach to problem solving is predicated on the Wisdom of Crowds concept that holds that under certain conditions the aggregation of independent knowledge-based solutions will outperform individual solutions. Improved performance using Wisdom of Crowds over individual solutions has been shown for classical search problems such as the traveling salesman problem given both human crowds (Yi, Steyvers, Lee, & Dry, 2012) and artificial crowds (Yampolskiy & Barkouky, 2011) and in problems in the domain of molecular biology such as inferring gene networks by combining the solutions of multiple experiments and models (Marbach et al., 2012). Wisdom of Crowds aggregation is fundamentally related to bootstrap aggregation or “bagging” (Hastie, Tibshirani, & Friedman, 2009), however we use the term Wisdom of Crowds to emphasize that consensus wisdom is applied to variable selection as a search problem as opposed to the aggregation of predictions. We loosely follow the process utilized by Yampolskiy and Barkouky (2011) in that we first generate individual solutions using a genetic algorithm and then aggregate a proportion of the solutions as “experts” to determine a consensus solution to variable selection. We then generate a classifier from the WoAC selected metabolites and evaluate classifier performance relative to classifiers constructed without WoAC that employ embedded feature selection (Elastic Net and Lasso multinomial classifiers).

## 2. Materials and Methods

### 2.1 Study Cohort

Patients presenting to two hospitals in Louisville, Kentucky with suspected acute myocardial infarction were enrolled in the study upon providing written informed consent. Additionally, patients presenting for an elective outpatient procedure for the treatment of stable coronary artery disease were enrolled as stable disease controls upon providing written informed consent. Novel criteria was developed by our group for differentiating thrombotic MI and non-thrombotic MI and has been previously discussed (DeFilippis et al., 2015) and is presented in Supplementary Table 1. Briefly, this criteria required clinical presentation consistent with the universal definition of AMI and a finding of positive Troponin for inclusion in either AMI study group. For the thrombotic MI group, recovery of a histologically confirmed coronary thrombus as well as ≥ 50% stenosis in the same vessel was an inclusion criteria. For inclusion in the non-thrombotic MI group, subjects must not have had significant stenosis or evidence of flow-limiting stenotic lesions evaluated by angiography and must not have had a thrombus recovered. The strict inclusion criteria was designed to limit misclassification of thrombotic MI and non-thrombotic MI subjects, and many borderline cases were eliminated (Supplemental Figure 2). The cohort described in this study included 11 thrombotic MI, 12 non-thrombotic MI, and 15 stable CAD subjects.

### 2.2 Plasma Samples and Analytical Measurement

Whole blood was collected from study subjects immediately prior to cardiac catheterization and plasma was extracted via centrifugation. The detection and quantification of metabolite relative abundances was conducted by Metabolon, Inc. Samples were analyzed by UPLC-MS/MS with positive ion mode electrospray ionization, negative ion mode electrospray ionization, and a negative ionization optimized for polar molecule detection and GC-MS. The identity of biochemicals detected was based on retention index matching, mass to charge ratio matching, and spectral data from libraries of known standards. All metabolites were identified at MSI level 1 unless denoted “unknown”. After identification, relative abundances were quantified by determining the area-under-the curve. After quantification, minimum values were imputed for missing abundances predicated on the assumption that these compounds were either not present or below the limit of detection. The resulting data was then median scaled and log-transformed (base 2).

### 2.3 Genetic Algorithm

We begin our description of the approach used to develop a classifier by introducing our notation. The phenotype (thrombotic MI, non-thrombotic MI, or stable CAD) of the *i*th plasma sample for *i* = 1,2, …, *N*, is represented as:

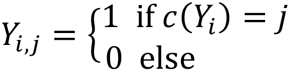

where *c*(*Y*_*i*_) returns a phenotype index and *j* ∈ {1,2, …, *J*} is the index of phenotypes. The vector **x**_*i*_ represents the metabolite abundances from the *i*th sample but the notation *X*_*m*_ is used to emphasize that the abundance of the *m*th metabolite is a random variable. The probability the *i*th sample has phenotype index *j* is then:

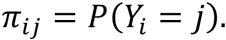

A multinomial logit model was assumed for determining the phenotype probabilities of each sample. This model is a generalized linear model with the following form (Agresti, 2013):

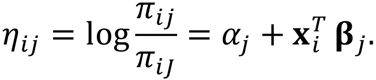

The estimated phenotype probabilities are then:

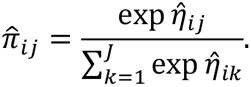

The subset of metabolites included as predictors in the multinomial model was denoted *M*, with the complement subset (metabolites not included) denoted *M*^c^.

To employ a Wisdom of Crowds approach for metabolite selection, a synthetic crowd was generated. A crowd was generated by first determining optimal metabolite subsets using a non-standard genetic algorithm. This algorithm emulated four biologically-inspired processes (birth/external immigration, recombination, mutation, and death) mimicking the evolution of chromosomes as the material of trait inheritance (Griffiths, Wessler, Carroll, & Doebley, 2015). Each iteration of the algorithm represented one temporal generation for the population of genetic material. The algorithm was initialized by generating an initial list of Boolean vectors *B*^(1)^ = {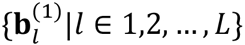 |*l* ∈ 1,2, …, *L*} with entries 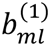 defined as:

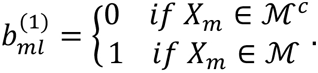

The initial Boolean vectors were generated by simulating Bernoulli random variables to generate a population with limiting distribution 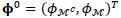

After initialization, the cost of each vector was estimated prediction error using repeated *k*-fold cross-validation. Each multinomial logit model was used to generate phenotype probability estimates *π̂*_*ij*_ From these estimates the cross-entropy loss (Bishop, 2006) was determined as:

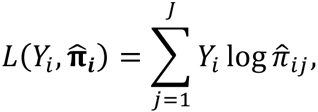

with corresponding cross-entropy error of prediction:

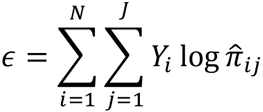

Cross-validation was used to estimate the expected error of prediction (Hastie et al., 2009). The observed data {(**x**_*i*_,*Y*_i_): *i* = 1,2, …, *N*} was randomly partitioned into *k* folds (we choose *k* = 10). Representing this partition as a mapping of samples to folds *κ*: {1,2, …, *N*} ↦ {1,2, …, *k*}, the cross-validation estimated error of prediction was then:

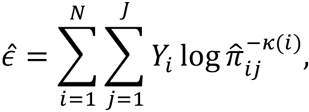

where 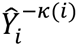 denotes the predicted phenotype of the *i*th sample with the *κ*(*i*) fold removed in the estimation of the multinomial logit model. Given the potential variability of cross-validation estimates of expected prediction error over small samples (Bengio & Grandvalet, 2004), repeated cross-validation was employed in which the random partitioning and estimation were conducted repeatedly (10 times) and the mean of the estimates was taken as the cost of the vector **b**_*l*_ The fitness of **b**_*l*_ was inversely proportional to the cost.

After determining the fitness of each inclusion vector, a two-point crossover recombination operator was used to generate child chromosomes (Nomura, 1997). This operator selected two chiasmata (location of crossover) two mimic a double crossover of homologous chromosomes (Griffiths et al., 2015). As a concrete example consider **b**_1_ = (0,1,0,1,0,1,0) and **b**_2_ = (1,0,1,0,1,0,1) for crossover with chiasmata at positions 2 and 5. Possible progeny vectors are then: **b**_3_ = (0,1|1,0,1|1,0) and **b**_4_ = (1,0|0,1,0|0,1). If a single endpoint was selected as chiasma, then a single-point crossover could be produced. The fitness of each **b**_*l*_ determined the probability that that vector would be a parent in recombination. The probability of recombination participation was determined by mapping the empirical cumulative distribution function (ECDF) of the fitness of each 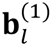to the cumulative distribution function of a Beta distribution, *Beta*(1,3), to generate Bernoulli priors, *p*_*l*_, where *p*_*l*_ was the probability of recombination participation. In addition to generating new genetic material via a recombination operator, new genetic material was generated by an “immigration” which created a small proportion of new inclusion vectors using the same generation function that generated the initial population. A death operator was defined for maintaining a stable population size and to retire inclusion vectors of low fitness. The death operator proceeded as follows: a cost function was defined as a weighted average of inclusion vector fitness (the inverse of repeated cross-validation estimated misclassification error) and age (the number of generations over which the vector had existed). As the recombination operator generated new genetic material, low fitness inclusion vectors were removed from the population by the death operator. A mutation operator was defined by generating a transition matrix **A** for each vector 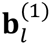∈ *B*^(1)^ to maintain the steady state distribution **π**^0^ = (π_*M*_*c*, *π*_*M*_)^*T*^, that is **Aπ**^(*t*)^ = **π**^0^ (although the mutation operator was applied only once per generation). The entries of the transition matrix were determined by varying the proportion of metabolites switching state inversely with fitness, that is the mutation rate φ_*l*_ = *a*_*M*,*M*_*c* + *a*_*M*_*c*,_*M*_ varied linearly with inverse fitness. Each binary metabolite indicator random variable 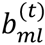thus was a discrete nonhomogeneous Markov chain (Lawler, 2006).

### 2.4 Artificial Crowds Aggregation

Following the method introduced in Yampolskiy and Barkouky (2011), we generate artificial crowd wisdom by aggregating intermediate genetic algorithm solutions. For crowd wisdom to generate “wise” solutions a crowd must be diverse (solutions determined from private information) and independent (Surowiecki, 2004). Similarly the aggregation of classifiers benefits from the introduction of diversity, such as with Random Forest ensembles (Breiman, 2001). As with Random Forest ensembles, we introduce diversity via bootstrapping and a random subspaces constraint. Specifically, 1,000 bootstrapped samples, (**X**^∗^, **Y**^∗^)_*k*_, were drawn with replacement from the original data. The columns of 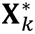 were then sampled without replacement to generate a new matrix 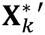 such that dim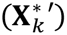 = *n*×floor(*p*/3). Each sample (**X**^∗^′, **Y**^∗^)_*k*_ was then used to initialize a genetic algorithm population. A single iteration of the algorithm consisted of the application of the following operators to the population: recombination, immigration, death, and mutation. Each initial population of inclusion vectors *B*^(1)^ had a population size of 250 and underwent 200 iterations, representing 200 generations of evolution. From each population of 250 inclusion vectors, the best solution of the inclusion vectors was retained as an “expert”. The proportion of times a metabolite was included in a multinomial model was then computed over the independent populations as: 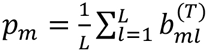 where *l* = 1,2, …, *L* indexed the independent GA populations.

The proportion *p*_*m*_ was then used as a measure of relative variable importance as it represented the propensity of a metabolite *m* to be included in an expert solution generated by genetic algorithm evolution. Metabolites that met a threshold of *p*_*m*_ > 0.025 were included in the final classifier. Given the objective of achieving maximal parsimony without compromising classification performance, the final multinomial logit model was fit via penalized maximum likelihood. The elastic-net penalty was used which includes *L*_1_ and *L*_2_ norm penalties (Zou & Hastie, 2005). Using the notation supplied of (Hastie et al., 2009), maximizing the penalized likelihood for a multinomial model with an elastic-net penalty corresponds to optimization of the following:

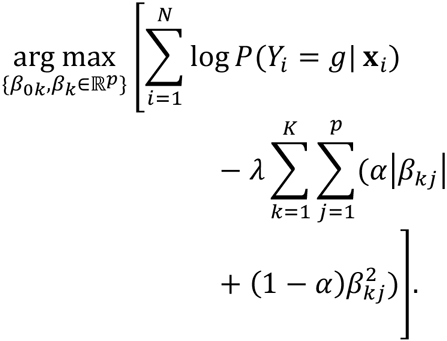

In this notation, *K* represents the number of phenotypes and *p* represents the number of metabolites in the model. The parameter α controls the tradeoff between the Lasso penalty and the ridge regression penalty.

### 2.4 Performance Evaluation

The performance of the classifier employing WoAC feature selection was evaluated by comparison with elastic net and Lasso (Tibshirani, 1996) regularized multinomial classifiers without WoAC feature selection (no-WoAC). The primary objective measure of classification performance was the repeated cross-validation estimated misclassification error rate. The number of metabolites selected by each method was also evaluated.

## 3. Results

The cost paths of the genetic algorithm (GA) solution exhibiting minimal final cost (cross-entropy loss) for 10 randomly selected populations are shown in Figure 1. These solutions are representative members of the Artificial Crowds. Diminishing returns in cross-entropy loss reduction are observed with increasing evolutionary time. The empirical distribution of *p*_*m*_ for the metabolites with a non-zero *p*_*m*_ value is shown in Figure 2. Of the 1,032 metabolites, 29 met the *p*_*m*_ criteria for variable selection. The selected metabolites consisted of: 5 lysophospholipids (LysoPE and LysoPC species), 5 steroid metabolites (pregnenolone and corticosteroid metabolites), 4 monoacylglycerols, 3 were amino acids (His, Lys, Ser), 3-aminoisobutyrate, 3-hydroxypyridine sulfate, 3-methyl catechol sulfate, 4-allylphenol sulfate, methyl-4-hydroxybenzoate, nicotinamide, phosphate, and 5 unknowns. The pairwise correlations between these metabolites are presented in Figure 3 and the abundance distributions of the selected metabolites are shown in Supplemental Figure 2. Significant correlations were observed in the steroid hormone abundances and in the monoacylglycerols abundances. The abundance of steroid hormones was negatively correlated with the abundance of Ser, Lys, His, nicotinamide, and phosphate. While not the primary evaluation, unsupervised hierarchical clustering showed minimal phenotype confusion given the abundances of metabolites selected by WoAC (Figure 4).

**Figure 1:**
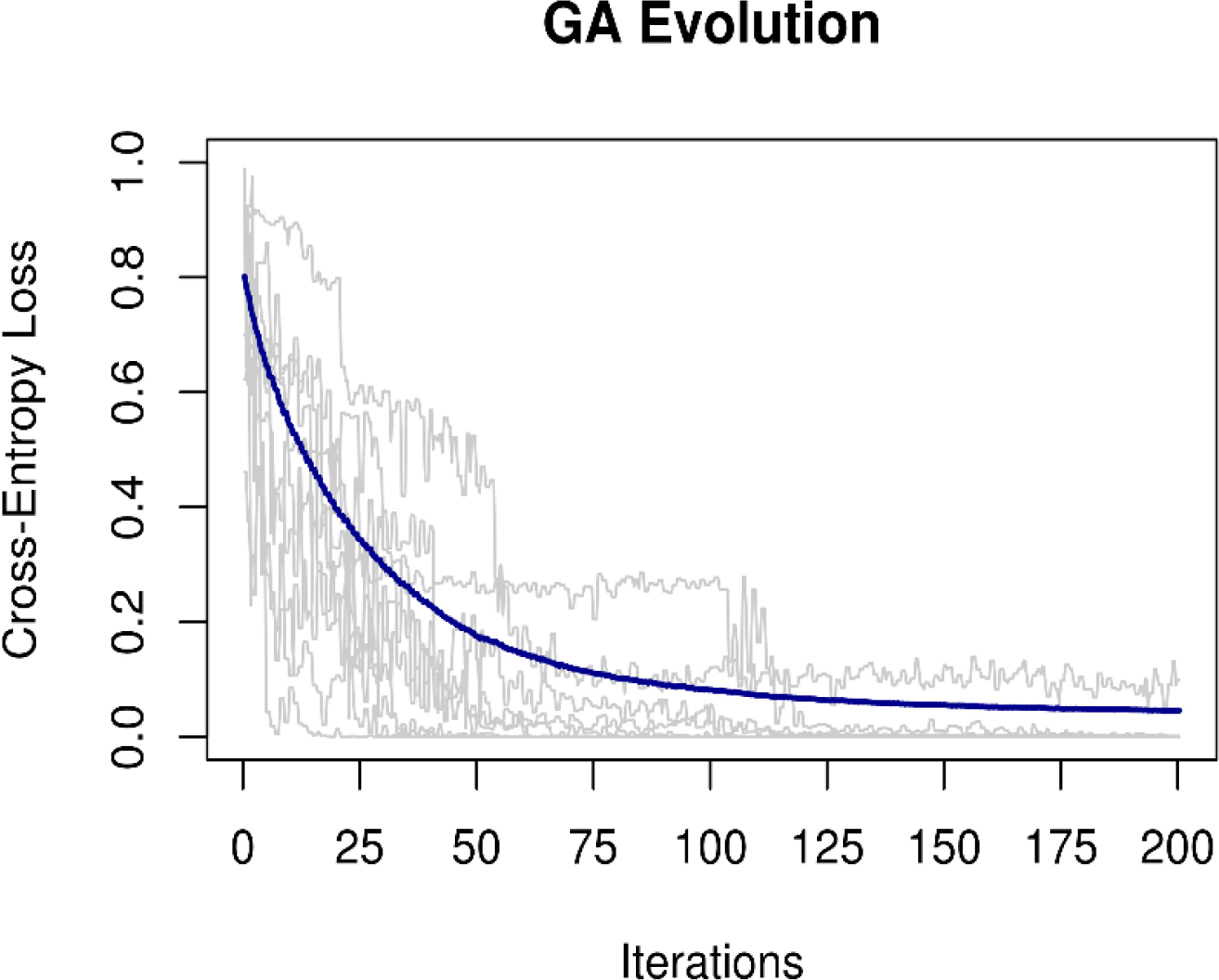
Cost-paths of the GA solution with the minimum final cost (cross-entropy loss) from 10 randomly selected populations. The average cost over all populations is also shown (blue line).

**Figure 2:**
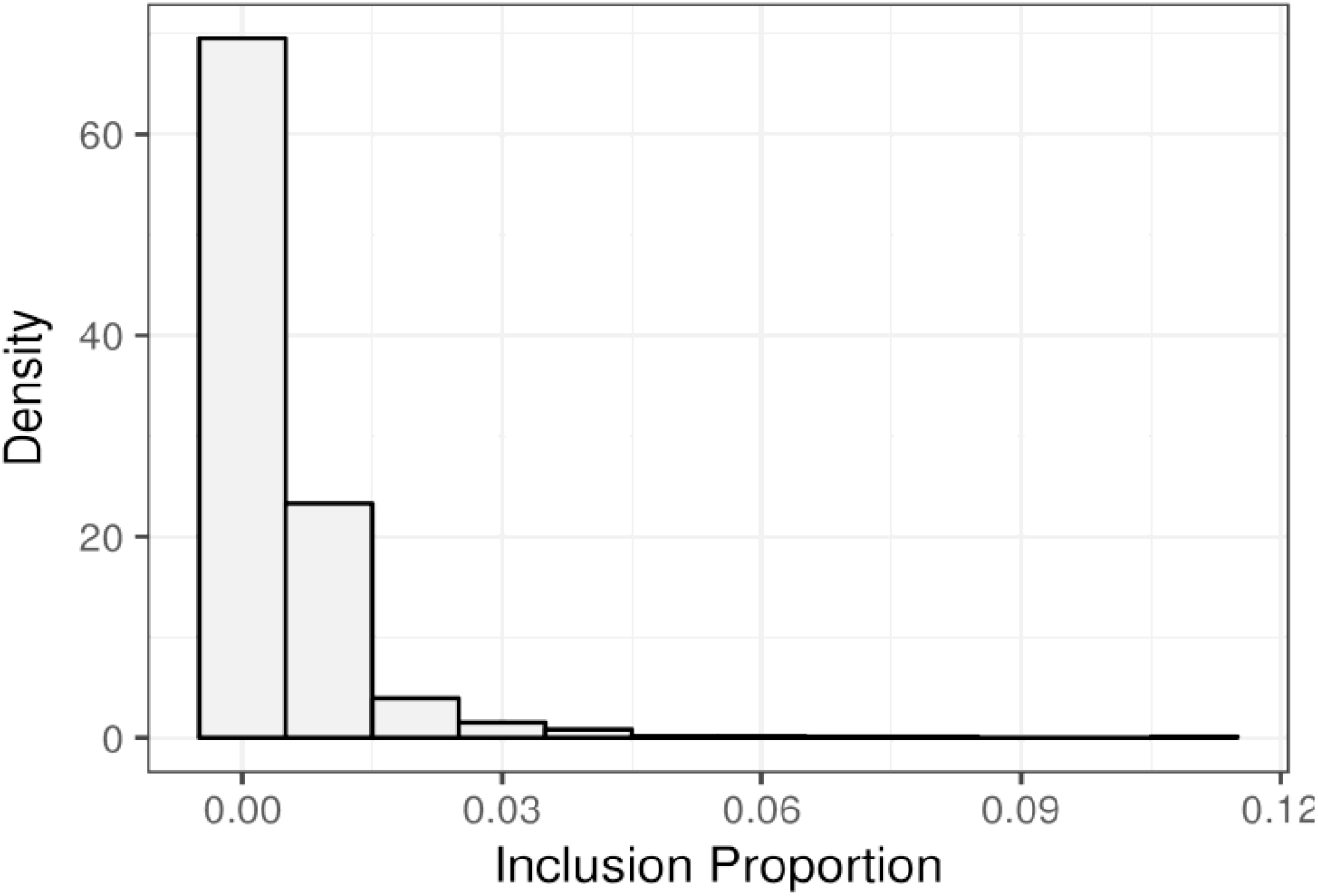
Empirical distribution of inclusion proportion for metabolites with p_m_ > 0.

**Figure 3:**
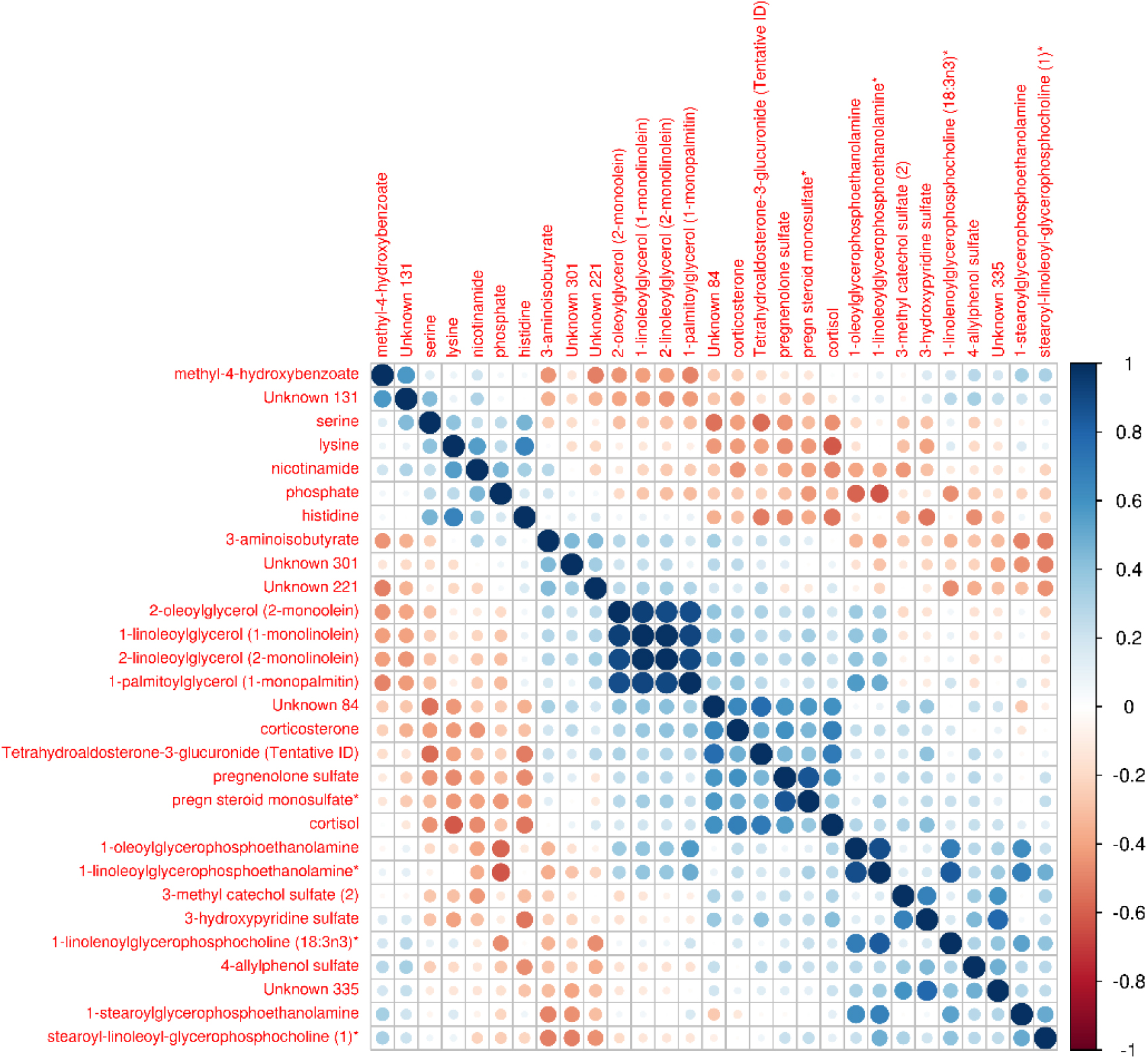
Pairwise correlations between WoAC selected metabolites

**Figure 4:**
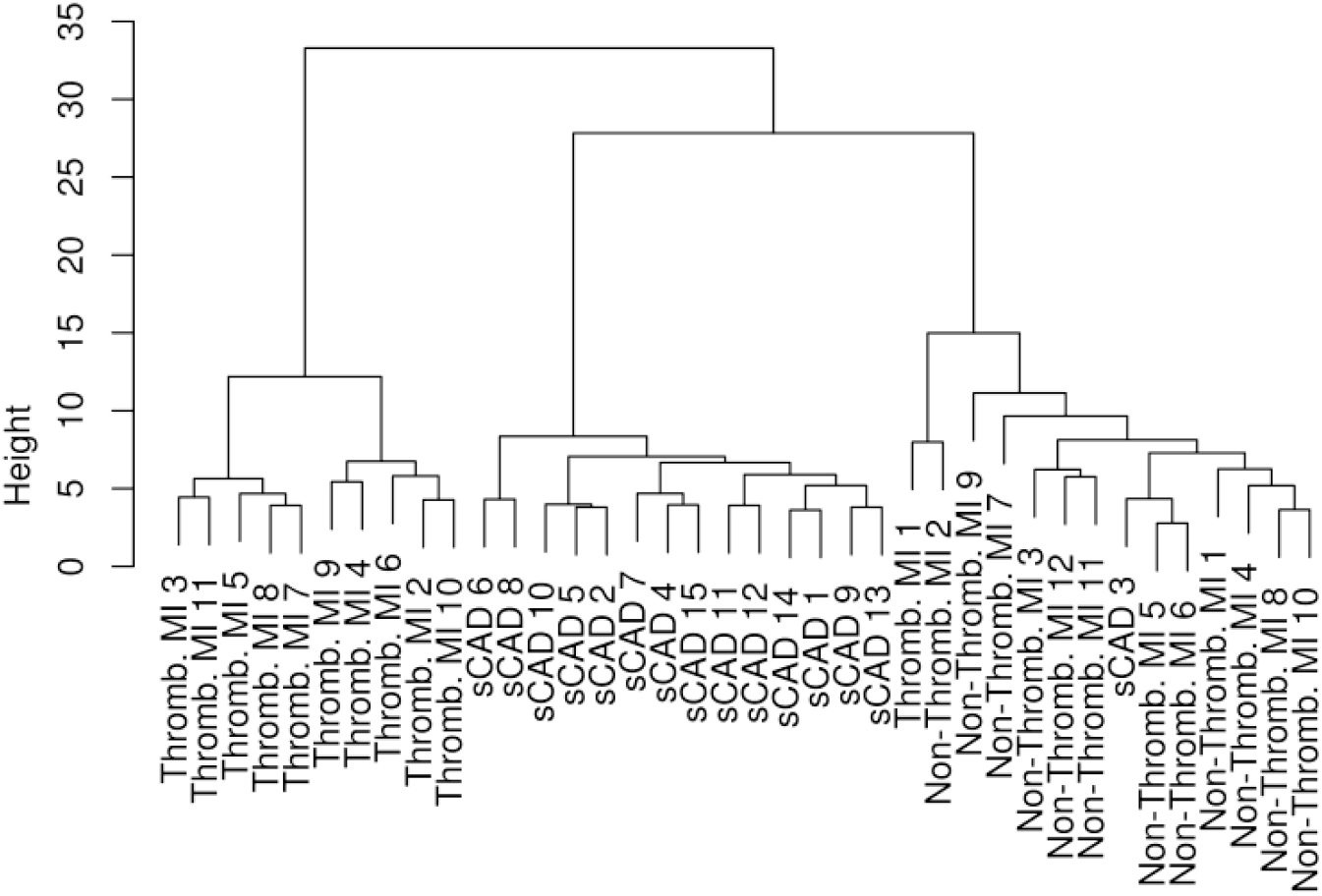
Hierarchical clustering of plasma samples using the metabolites selected by WoAC

The results of the evaluation of WoAC variable selection performance are presented in Table 1 and Figure 5. For each value of α, the parameter that controls the weighting of *L*_1_ and *L*_2_ norm penalties, employing WoAC variable selection led to a lower estimated misclassification rate. In employing the elastic net a trade-off between model accuracy and shrinkage was observed with the manipulation of α. The benefit of WoAC feature selection prior to classifier construction when shrinkage is prioritized can be seen in the Lasso case. In this case, the number of metabolites selected was similar between the WoAC and no-WoAC conditions, with an average of 8.3 and 9.1 metabolites selected, respectively; the median estimated misclassification rate was approximately halved from 26.3% to 13.2% with the employment of WoAC. The minimum estimated misclassification rate achieved in the WoAC and no-WoAC conditions was 2.6% and 18.4% with α = .25, with an average of 16.4 and 68.0 metabolites included in the models, respectively.

**Table 1:** Misclassification rate and number of metabolites selected by method. Misclassification rate and number selected were estimated via repeated cross-validation.

| Alpha | Misclassification Rate (Median, IQR) |  | Metabolites Selected (Mean, SD) |  |
| --- | --- | --- | --- | --- |
|  | WoAC | No WoAC | WoAC | No WoAC |
| .25 | .026, .000 | .184, .053 | 16.4, 1.1 | 68.0, 14.4 |
| .50 | .053, .026 | .237, .053 | 13.1, 1.2 | 36.0, 10.1 |
| .75 | .079, .026 | .263, .053 | 11.6, 1.7 | 16.4, 5.9 |
| Lasso | .132, .053 | .263, .079 | 8.3, 1.0 | 9.1, 1.7 |

**Figure 5:**
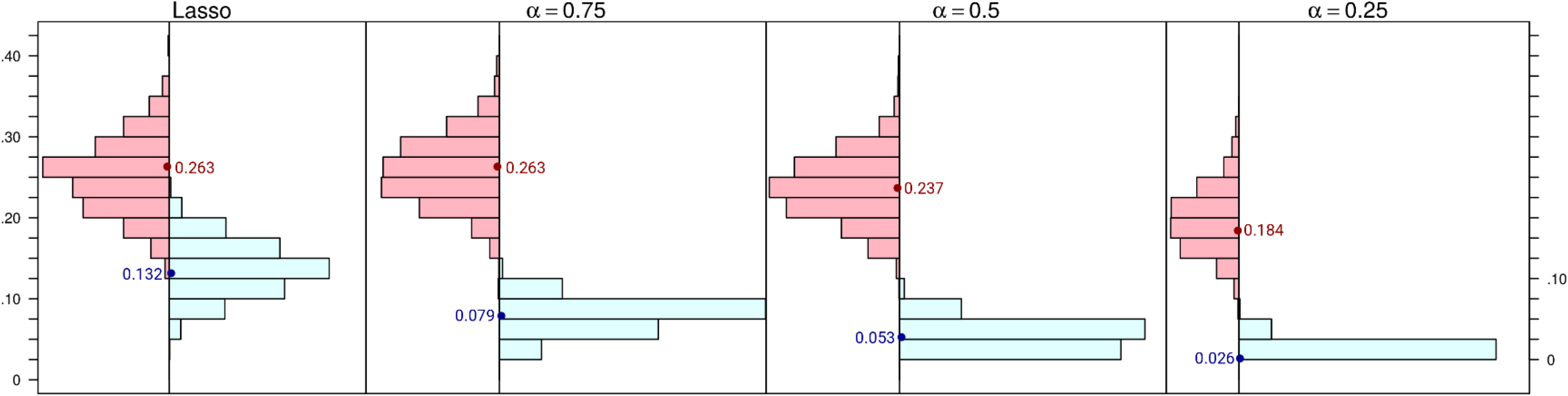
Repeated cross-validation estimated misclassification rate distribution for lasso and elastic net multinomial classifiers with WoAC metabolite selection (blue) and without (red). 100 replications of 10-fold CV were used to estimate error. Median of each distribution is shown as a dot.

## 4. Discussion

In this paper we detailed an evaluation of Wisdom of Artificial Crowds using intermediate genetic algorithm solutions for variable selection. The successful application of genetic algorithms to feature selection in metabolomics is well established with early papers pairing GA variable selection with partial least squares regression (PLSR) for determining important spectral features from mass spectrometry (Broadhurst, Goodacre, Jones, Rowland, & Kell, 1997) and nuclear magnetic resonance (Leardi & Lupiáñez González, 1998) data. However, unlike these works, we do not directly incorporate a genetic algorithm solution or set of solutions directly via embedding. Instead, our application first generates independently evolved and diverse (via bootstrapping and the random subspaces constraint) populations of genetic algorithm solutions as a crowd and extracts the consensus wisdom of that crowd. This wisdom is then applied to the variable selection problem, which then enables improved classifier construction.

In the classifier construction, variable selection was again carried out via penalized maximum likelihood estimation. We observed that using the elastic net that a lower ratio of *L*_2_ to *L*_1_ norm penalization generated a better result following WoAC variable selection. This is an aspect that deserves further exploration in future work. Multidimensional cross-validation could be used to determine the optimal parameter values for both *p*_*m*_, which controls the number of metabolites selected via WoAC, and α which controls the shrinkage of model parameters. A significant challenge in this effort would be determining a suitable cost function. The loss associated with misclassification and the loss associated with increasing the number of metabolites must be combined into a cohesive cost function. Further, differential penalization of misclassification by phenotype may be advisable based on clinical consequence.

The variables selected by WoAC report some of the metabolic consequences of acute thrombotic MI. Specifically, an increased abundance of selected pregnenolone and corticosteroid metabolites was observed in the thrombotic MI group relative to the non-thrombotic and stable CAD groups. This may be indicative of stimulation of the hypothalamic-pituitary-adrenal axis following thrombotic MI. Evidence of activation of this axis following MI has been demonstrated in other studies that have shown increased levels of circulating adrenocorticotropic hormone (Paganelli et al., 2003) and copeptin (Maisel et al., 2013) in the hours following acute MI. As this signal was stronger in the thrombotic MI group than the non-thrombotic MI group (especially for cortisol), an increase in these hormones in circulation may be associated with thrombosis directly in addition to acute stress. Others have demonstrated a mechanistic relationship between platelet activating factor, an important factor in stimulating platelet activation and aggregation, and glucocorticoids (Aikawa et al., 1991; Shimada, Hirose, Matsumoto, & Aikawa, 2005). Increased abundance of selected monoacylglycerols was observed in both acute MI groups relative to the stable CAD group and these abundance distributions exhibited significant pairwise correlations. An increased abundance of monoacylglycerols may be indicative of increased hydrolysis of triacylglycerol molecules or impaired uptake of these molecules from plasma. Decreased plasma concentrations of selected amino acids in the thrombotic MI group relative to the non-thrombotic and stable CAD groups is an interesting finding and may indicate increased catabolism of amino acids to furnish ATP for the ischemic heart. All of the amino acids identified can be catabolized to produce ATP either via gluconeogenesis or ketogenesis (Voet, Voet, & Pratt, 2013). Under ischemic conditions, the heart must utilize metabolic substrates that do not require oxygen (Drake, Sidorov, McGuinness, Wasserman, & Wikswo, 2012); hence, the inability to oxidize fatty acids may lead to increased utilization of amino acids. Alternatively, a decrease in plasma amino acid concentrations may be due to increased protein synthesis in activated platelets. Platelet activation results in signal dependent translation of factors involved in thrombosis such as tissue factor (TF) (Panes et al., 2007) and plasminogen activator inhibitor-1 (PAI-1) (Brogren, 2004), which could result in diminished concentrations of amino acids following MI in thrombotic MI subjects.

The most significant finding of this work is that applying WoAC feature selection prior to classifier construction significantly improved classifier accuracy relative to regularization methods for our application. The application of WoAC feature selection with an elastic net multinomial regression classifier furnished a diagnostic classifier that demonstrated minimal estimated misclassification error (2.6%) and high estimated sensitivity (90.9%) and specificity (100.0%) for detecting thrombotic MI. Given the potential clinical relevance of such a blood-based test, this classifier warrants further validation in an independent cohort.

## Acknowledgements

The authors would like to thank the human participants who graciously agreed to take part in the study. The authors thank the members of the Atherosclerosis and Atherothrombosis Research laboratory at the University of Louisville for help with sample processing and Samantha Carlisle, M.S. for her assistance.

---

**Supplemental Figure 1:**
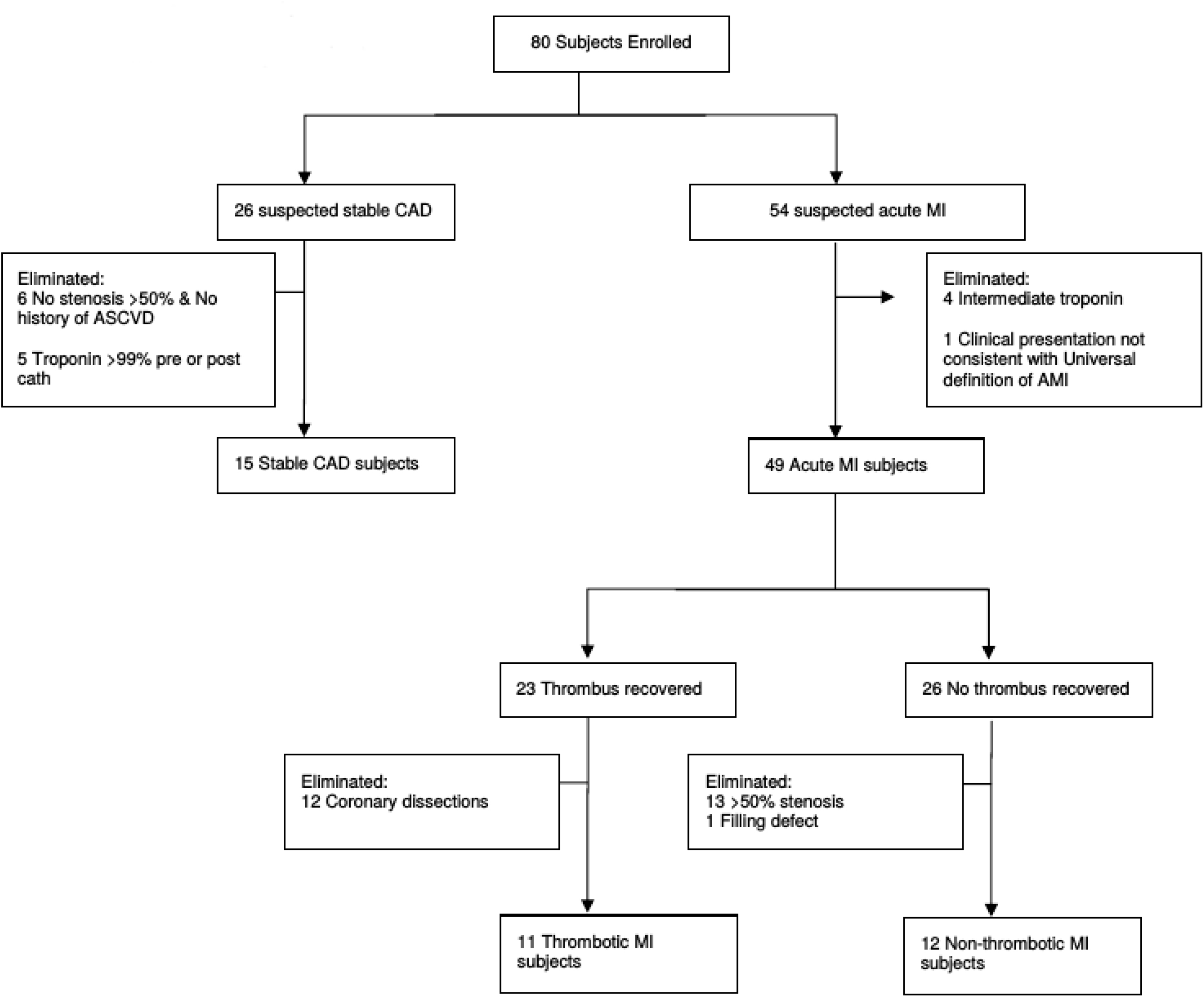
Flowchart depicting enrollment inclusion and exclusion criteria for the clinical cohort.

**Supplemental Figure 2:**
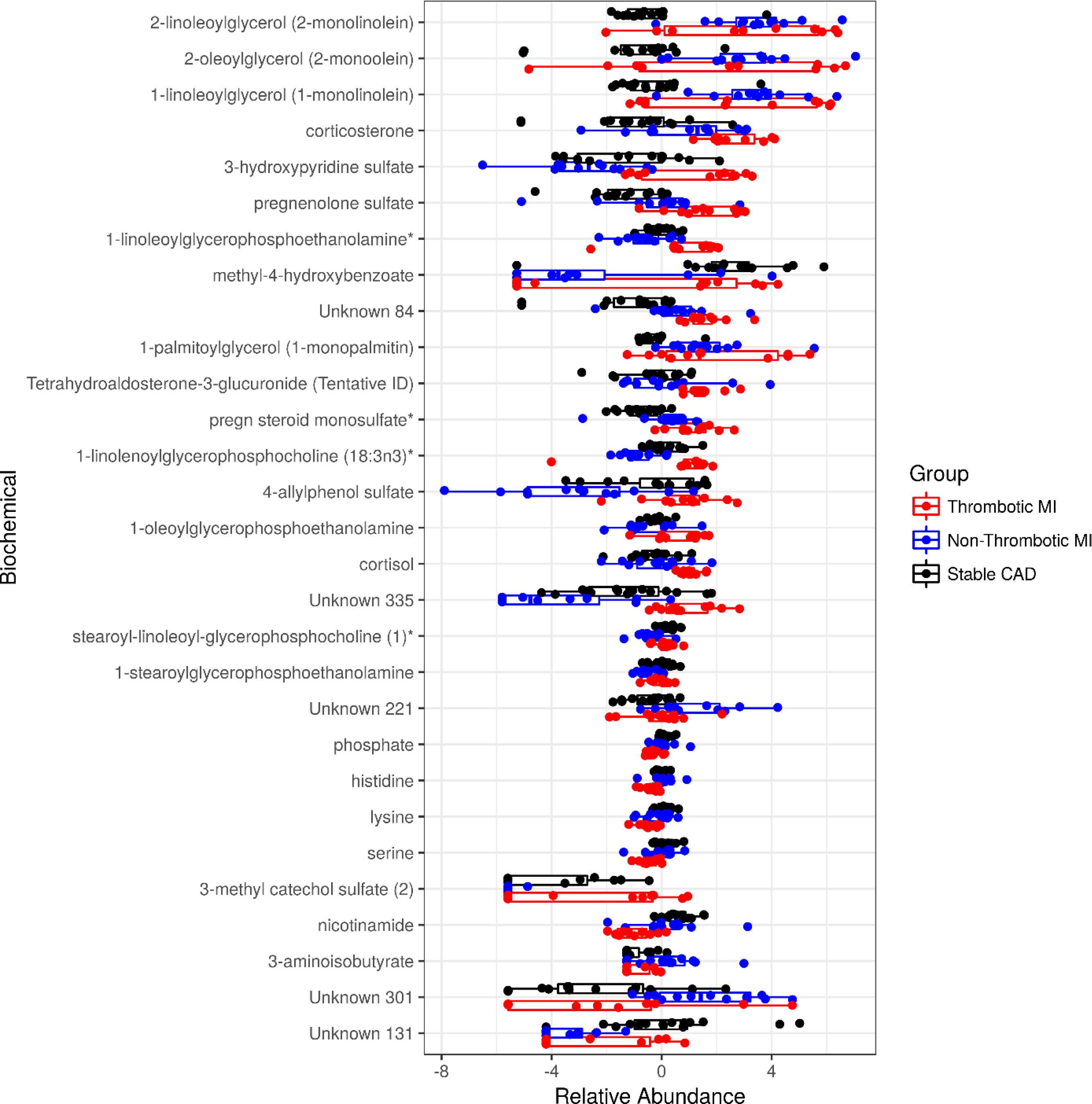
WoAC selected metabolite abundances by group. Log-transformed median-scaled relative abundances are shown for each metabolite.

## References

Agresti, A. (2013). Categorical data analysis (3rd ed.). Hoboken, NJ: Wiley.

Aikawa, T., Hirose, T., Matsumoto, I., Morikawa, T., Shimada, T., Mine, Y., … Tsuji, Y. (1991). Effect of platelet-activating factor on cortisol and corticosterone secretion by perfused dog adrenal. Lipids, 26(12), 1108-1111.

Amsterdam, E. A., Wenger, N. K., Brindis, R. G., Casey, D. E., Jr., Ganiats, T. G., Holmes, D. R., Jr., … American Association for Clinical, C. (2014). 2014 AHA/ACC Guideline for the Management of Patients with Non-ST-Elevation Acute Coronary Syndromes: a report of the American College of Cardiology/American Heart Association Task Force on Practice Guidelines. J Am Coll Cardiol, 64(24), e139-228. doi:10.1016/j.jacc.2014.09.017

Bengio, Y., & Grandvalet, Y. (2004). No unbiased estimator of the variance of k-fold cross-validation. Journal of Machine Learning Research, 5(Sep), 1089-1105.

Bishop, C. M. (2006). Pattern recognition and machine learning. New York: Springer.

Breiman, L. (2001). Random Forests. Machine Learning, 45(1), 5-32. doi:10.1023/a:1010933404324

Broadhurst, D., Goodacre, R., Jones, A., Rowland, J. J., & Kell, D. B. (1997). Genetic algorithms as a method for variable selection in multiple linear regression and partial least squares regression, with applications to pyrolysis mass spectrometry. Analytica Chimica Acta, 348(1-3), 71-86. doi:10.1016/s0003-2670(97)00065-2

Brogren, H. (2004). Platelets synthesize large amounts of active plasminogen activator inhibitor 1. Blood, 104(13), 3943-3948. doi:10.1182/blood-2004-04-1439

DeFilippis, A. P., Chernyavskiy, I., Amraotkar, A. R., Trainor, P. J., Kothari, S., Ismail, I., … Bhatnagar, A. (2015). Circulating levels of plasminogen and oxidized phospholipids bound to plasminogen distinguish between atherothrombotic and non-atherothrombotic myocardial infarction. J Thromb Thrombolysis. doi:10.1007/s11239-015-1292-5

Drake, K. J., Sidorov, V. Y., McGuinness, O. P., Wasserman, D. H., & Wikswo, J. P. (2012). Amino acids as metabolic substrates during cardiac ischemia. Exp Biol Med (Maywood), 237(12), 1369-1378. doi:10.1258/ebm.2012.012025

Griffiths, A. J. F., Wessler, S. R., Carroll, S. B., & Doebley, J. F. (2015). Introduction to genetic analysis (Eleventh edition. ed.). New York, NY: W.H. Freeman & Company, a Macmillan Education imprint.

Hastie, T., Tibshirani, R., & Friedman, J. H. (2009). The elements of statistical learning: data mining, inference, and prediction (2nd ed.). New York, NY: Springer.

Lawler, G. F. (2006). Introduction to stochastic processes (2nd ed.). Boca Raton: Chapman & Hall/CRC.

Leardi, R., & Lupiáñez González, A. (1998). Genetic algorithms applied to feature selection in PLS regression: how and when to use them. Chemometrics and Intelligent Laboratory Systems, 41(2), 195-207. doi:10.1016/s0169-7439(98)00051-3

Maisel, A., Mueller, C., Neath, S. X., Christenson, R. H., Morgenthaler, N. G., McCord, J., … Peacock, W. F. (2013). Copeptin helps in the early detection of patients with acute myocardial infarction: primary results of the CHOPIN trial (Copeptin Helps in the early detection Of Patients with acute myocardial INfarction). J Am Coll Cardiol, 62(2), 150-160. doi:10.1016/j.jacc.2013.04.011

Marbach, D., Costello, J. C., Küffner, R., Vega, N. M., Prill, R. J., Camacho, D. M., … Stolovitzky, G. (2012). Wisdom of crowds for robust gene network inference. Nature Methods, 9(8), 796-804. doi:10.1038/nmeth.2016

Mozaffarian, D., Benjamin, E. J., Go, A. S., Arnett, D. K., Blaha, M. J., Cushman, M., … Stroke Statistics, S. (2016). Heart Disease and Stroke Statistics-2016 Update: A Report From the American Heart Association. Circulation, 133(4), e38-e360. doi:10.1161/CIR.0000000000000350

Newby, L. K., Jesse, R. L., Babb, J. D., Christenson, R. H., De Fer, T. M., Diamond, G. A., … Wesley, D. J. (2012). ACCF 2012 Expert Consensus Document on Practical Clinical Considerations in the Interpretation of Troponin Elevations. Journal of the American College of Cardiology, 60(23), 2427-2463. doi:10.1016/j.jacc.2012.08.969

Nicholson, J. K., Holmes, E., Kinross, J. M., Darzi, A. W., Takats, Z., & Lindon, J. C. (2012). Metabolic phenotyping in clinical and surgical environments. Nature, 491(7424), 384-392. doi:10.1038/nature11708

Nomura, T. (1997). An analysis on linear crossover for real number chromosomes in an infinite population size. 111-114. doi:10.1109/icec.1997.592279

Paganelli, F., Frachebois, C., Velut, J. G., Boullu, S., Sauze, N., Rosso, J. P., … Oliver, C. (2003). Hypothalamo-pituitary-adrenal axis in acute myocardial infarction treated by percutaneous transluminal coronary angioplasty: effect of time of presentation. J Endocrinol Invest, 26(5), 407-413. doi:10.1007/BF03345195

Panes, O., Matus, V., Saez, C. G., Quiroga, T., Pereira, J., & Mezzano, D. (2007). Human platelets synthesize and express functional tissue factor. Blood, 109(12), 5242-5250. doi:10.1182/blood-2006-06-030619

Psychogios, N., Hau, D. D., Peng, J., Guo, A. C., Mandal, R., Bouatra, S., … Wishart, D. S. (2011). The human serum metabolome. PLoS One, 6(2), e16957. doi:10.1371/journal.pone.0016957

Shimada, T., Hirose, T., Matsumoto, I., & Aikawa, T. (2005). Platelet-activating factor acts on cortisol secretion by perfused guinea-pig adrenals via calcium-/phospholipid-dependent mechanisms. J Endocrinol, 184(2), 381-391. doi:10.1677/joe.1.05937

Surowiecki, J. (2004). The wisdom of crowds: why the many are smarter than the few and how collective wisdom shapes business, economies, societies, and nations (1st ed.). New York: Doubleday:.

Thygesen, K., Alpert, J. S., Jaffe, A. S., Simoons, M. L., Chaitman, B. R., White, H. D., … Wagner, D. R. (2012). Third universal definition of myocardial infarction. J Am Coll Cardiol, 60(16), 1581-1598. doi:10.1016/j.jacc.2012.08.001

Tibshirani, R. (1996). Regression Shrinkage and Selection via the Lasso. Journal of the Royal Statistical Society. Series B 58(1).

Voet, D., Voet, J. G., & Pratt, C. W. (2013). Fundamentals of biochemistry: life at the molecular level (4th ed.). Hoboken, NJ: Wiley.

Yampolskiy, R. V., & Barkouky, A. E. L. (2011). Wisdom of artificial crowds algorithm for solving NP-hard problems. International Journal of Bio-Inspired Computation, 3(6), 358. doi:10.1504/ijbic.2011.043624

Yi, S. K., Steyvers, M., Lee, M. D., & Dry, M. J. (2012). The wisdom of the crowd in combinatorial problems. Cogn Sci, 36(3), 452-470. doi:10.1111/j.1551-6709.2011.01223.x

Zou, H., & Hastie, T. (2005). Regularization and variable selection via the elastic net. Journal of the Royal Statistical Society: Series B (Statistical Methodology), 67(2), 301-320. doi:10.1111/j.1467-9868.2005.00503.x

